# Phylogenomics, biogeography, and trait evolution of the Boletaceae (Boletales, Agaricomycetes, Basidiomycota)

**DOI:** 10.1101/2023.10.18.563010

**Authors:** Keaton Tremble, Terry Henkel, Alexander Bradshaw, Colin Domnauer, Lyda Brown, Lê Xuân Thám, Guliana Furci, Cathie Aime, Jean-Marc Moncalvo, Bryn Dentinger

## Abstract

The species-rich porcini mushroom family Boletaceae is a widespread and well-known group of ectomycorrhizal (ECM) mushroom-forming fungi that has eluded intrafamilial phylogenetic resolution despite many attempts using morphological traits and multi-locus molecular datasets. In this study, we present a genome-wide molecular dataset of 1764 single-copy gene families from a global sampling of 418 Boletaceae specimens. The resulting phylogenetic analysis has strong statistical support for most branches of the tree, including the first statistically robust backbone. The enigmatic *Phylloboletellus chloephorus* from non-ECM Argentinian subtropical forests was recovered as an early diverging lineage within the Boletaceae. Time-calibrated branch lengths estimate that the family first arose in the early- to mid-Cretaceous and underwent a rapid radiation in the Eocene, possibly when the ECM nutritional mode arose with the emergence and diversification of ECM angiosperms. Biogeographic reconstructions reveal a complex history of vicariance and episodic long-distance dispersal correlated with historical geologic events, including Gondwanan origins and cladogenesis patterns that parallel its fragmentation. Ancestral state reconstruction of sporocarp morphological traits predicts that the ancestor of the Boletaceae was lamellate with ornamented basidiospores, contrary to most contemporary “bolete” morphologies. Transition rates indicated that the lamellate hymenophore and sequestrate sporocarp are reversible traits. Together, this study represents the most comprehensively sampled, data-rich molecular phylogeny of the Boletaceae to date, enabling robust inferences of trait evolution and biogeography in the group.

## INTRODUCTION

Evolutionary radiations result from short bursts of relatively rapid diversification and demonstrate the creative power of evolution. Adaptive and non-adaptive radiations may be the most common macroevolutionary patterns and are fundamental to the origins of biodiversity (Simoes et al. 2016; Futuyma 1998; Schluter and McPeek 2000). Yet, the causes of evolutionary radiations are poorly known for most groups, as most studies have focused on animals and plants (Soulebeau et al. 2015). The other major multicellular eukaryotic group, the Fungi, have largely been neglected (Varga et al. 2019).

The porcini mushroom family Boletaceae (FIG. 1) is an example of an evolutionary radiation in Fungi (Bruns et al. 1992). The family is exceptionally diverse (>2000 currently accepted species) and globally distributed, but poorly documented for many regions. Yet, boletoid fungi are prevalent ectomycorrhizal (ECM) mutualists in ecosystems dominated by ECM plants (Peay et al. 2010) and at least eight species are traded globally as wild-collected edible mushrooms (Arora 2008; Sitta and Floriani 2008; Dentinger et al. 2010; Dentinger and Suz 2014). Despite their conspicuous sporocarps, ecological dominance, and cultural importance, new species of Boletaceae are regularly described from poorly explored habitats around the globe (e.g. Halling et al. 2006, 2023; Fulgenzi et al. 2007, 2008, 2010; Neves et al. 2010; Husbands et al. 2013; Castellano et al. 2016; Chakraborty et al. 2015; Das et al. 2015, 2016; Henkel et al. 2016; Magnago et al. 2017). New species have also recently been described from wild-collected foods in markets (e.g. Das et al. 2015; Dentinger and Suz 2014; Halling et al. 2014). While new Boletaceae species are increasingly understood in a global phylogenetic context, shedding light on their origin, diversification, and migration, over 20 years of molecular phylogenetic studies using legacy loci have made little progress towards resolving the deepest nodes (“backbones”) in Boletaceae phylogenies (Grubisha et al. 2001; Binder and Hibbett 2006; Drehmel et al. 2008; Dentinger et al. 2010; Nuhn et al. 2016; Wu et al. 2014).

**Figure 1.**
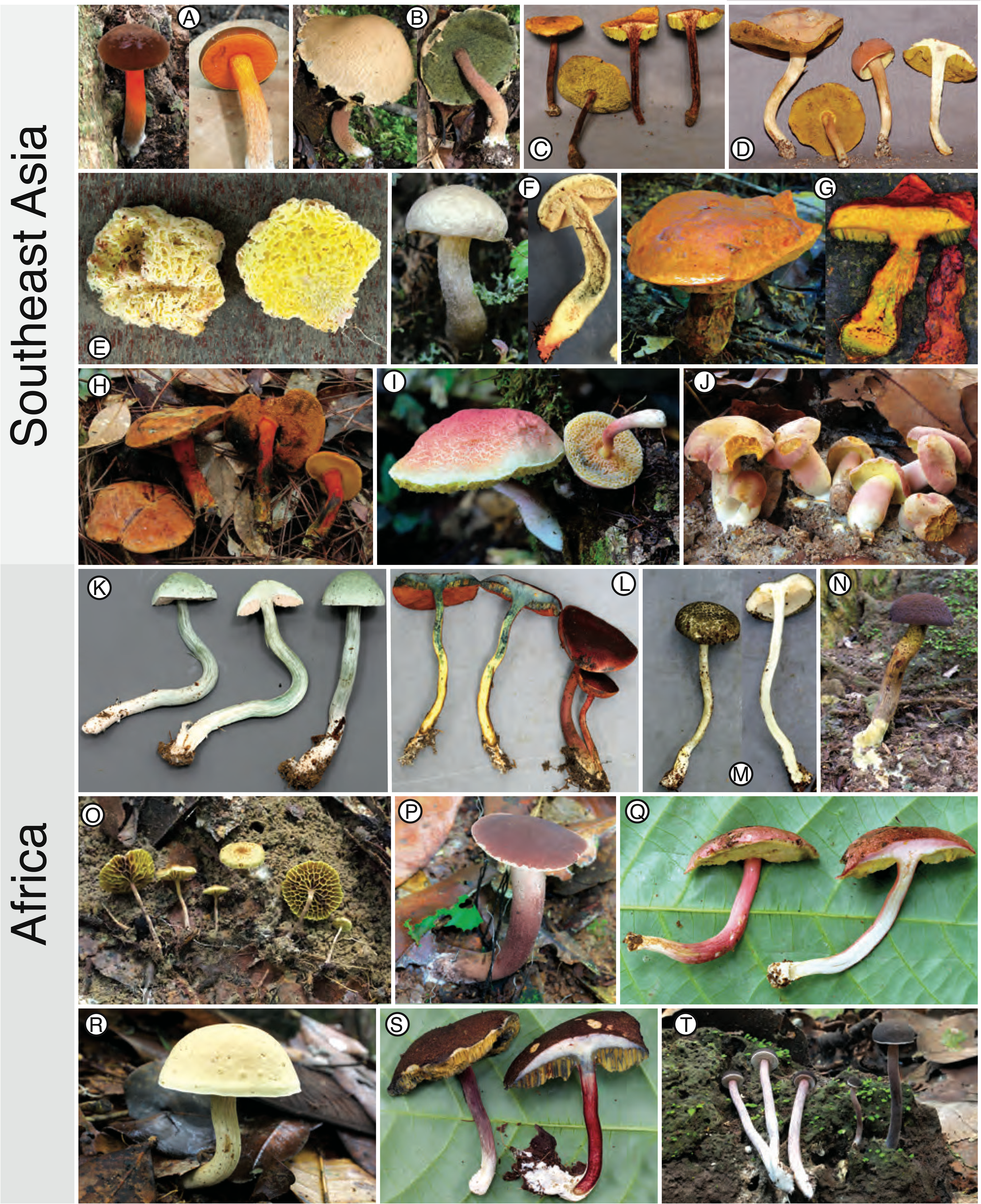
Selected Boletaceae collections from two of the most species-rich regions that were newly sequenced in this study. A) *Boletus cervinococcineus*, Singapore (BD616); B) *Heimioporus punctisporus*, Sarawak (BAKO2); C) unidenti-fied Boletaceae, Vietnam (CTN-08-0007); D) unidentified Boletaceae (CTN-08-0029); E) *Spongiforma* sp., Sarawak (BTNG10); F) *Leccinum* sp., Sarawak (SWK246); G) *Crocinoboletus laetissimus*, Sarawak (SWK335); H) unidentified Boletaceae sp., Viet-nam (DLT-08-0127); I) *Boletellus* sp., Sarawak (SWK356); J) unidentified Boletaceae, Vietnam (CTN-08-0051); K) *Tylopilus* sp., Cameroon (BD655); L) *Xerocomus* sp. 9. Cameroon (BD773); M) *Fistulinella staudtii*, Cameroon (BD848); N) *Boletellus* sp., Cameroon (BD714); O) *Phylloporus cf. tubipes*, Cameroon (BD719); P) *Tylopilus s*p. 8, Cameroon (BD816); Q) *Xerocomus* sp. 8, Cameroon (BD695); R) *Boletus alliaceus*, Cameroon (BD697); S) *Tubosaeta brunneosetosa*, Cameroon (BD686); T) *Tylopilus* sp., Cameroon (BD716). Not to scale.

A prominent consequence of this lack of phylogenetic resolution is the recent explosion of new generic names to accommodate newly discovered species, or species that are included in molecular phylogenetic analyses for the first time and recovered on long branches with no supported affinity to existing named genera (e.g., Castellano et al. 2016; Henkel et al. 2016; Badou et al. 2022; Halling et al. 2023). Few of these studies have followed recommended best practices for naming new genera (Vellinga et al. 2015). Moreover, many of these new Boletaceae genera are monotypic and require identification to recognize, an impractical solution to the problem. Taken together, the current situation is perhaps best described as a quagmire of nomenclatural, taxonomic, ecological, and evolutionary speculation.

Beyond taxonomic concerns, the Boletacaeae presents a unique system to identify the genetic mechanisms that contribute to diversification. The Boletaceae appear to have underwent an early evolutionary radiation between 60-100 mya (Bruns and Palmer 1989; Binder et al. 2006; Dentinger et al. 2010; Wu et al. 2014, 2016; Sato et al. 2019). This early radiation has been correlated with the convergent evolution of morphological traits, such as the lamellate hymenophore and gasteromycetization (Castellano et al. 2016), suggesting that emergence of morphological diversity is constrained by relatively few changes in developmental pathways.

Many factors have contributed to difficulties in generating robust phylogenetic reconstructions for the Boletaceae. While phenomena such as incomplete lineage sorting and hybridization may obscure historical phylogenetic signal, previous datasets for the Boletaceae had patchy taxonomic and geographic sampling. These factors impact accurate phylogenetic reconstruction, possibly exacerbated by the aforementioned rapid radiation event (Bruns et al. 1992; Sato et al. 2017). Without a phylogeny that is based on globally representative taxon sampling and statistically well-supported resolution at all depths of the tree, it is impossible to name, classify, and understand evolutionary history in the Boletaceae. For example, only a few recent studies have included representatives of the exceptionally rich Australian boletoid funga (Halling, Fechner, et al. 2015, 2023; Halling, Nuhn, et al. 2012). Boletoid fungi from the Neotropics and Afrotropics are rarely represented in family-level analyses despite their exceptional species richness (e.g. Heinemann 1951; Henkel et al. 2012).

Recent fieldwork has resulted in many new collections from under-sampled regions (B. Dentinger, T. Henkel, R. Halling; unpublished data), and these are now available to include in phylogenetic datasets aiming to achieve the first globally representative sampling of the Boletaceae. Collections-based phylogenomics is also effective for resolving ancient relationships among mushroom-forming fungi (Dentinger et al. 2015; Liimatainen et al. 2021; Tremble et al. 2020). However, no one has yet applied these methods to the Boletaceae. Moreover, whole genome sequencing of mushroom forming fungi provides opportunities to go beyond phylogenetic reconstruction. For example, whole genome sequencing can exceed legacy loci in identifying population processes that generate biodiversity (e.g. Tremble et al. 2022).

For this study, we generated the first phylogeny of the Boletaceae that utilized a very large molecular dataset comprised of 1764 genome-wide loci from 418 taxa across the family. Specimens were collected in many tropical and temperate geographic regions. To obtain broad geographic and taxon coverage we included type fungarium specimens and recent new collections. We utilized specimens from previously under-sampled regions including tropical Africa, southern South America, lowland tropical South America, and Australia. Type species were sampled to facilitate future taxonomic revisions of genera. Using our highly resolved genome-based phylogeny, we also performed the first inclusive biogeographic reconstruction of the Boletaceae and assessed morphological trait evolution. Overall, we provide new insights into the broad patterns of evolution of this enigmatic fungal group.

## MATERIALS AND METHODS

### Sampling

Taxon selection focused on obtaining representatives of all currently accepted genera, selecting type species whenever possible. Because the current understanding of genera is incomplete and rapidly changing, we could not include representatives of all currently accepted genera that were published during the course of this study. Specimens from geographic regions unrepresented in prior studies were also included. A total of 418 Boletaceae specimens were gathered from a global distribution using collections made by the authors, those borrowed from four institutions, and donations from citizen scientists (SUPPLEMENTAL TABLE 1). In addition, we utilized genome data publicly available from the JGI Mycocosm Portal (Grigoriev et al. 2014) for *Boletus coccyginus*, *B. reticuloceps*, *Butyriboletus roseoflavus*, *Chiua virens*, *Lanmaoa asiatica*, and *Imleria badia* (Miyauchi et al. 2020, Wu et al. 2022, Kohler et al. 2015). *Paxillus involutus*, *Paxillus adelphus*, and *Hydnomerulius pinastri* genomes from JGI were used for outgroups (Kohler et al. 2015).

### DNA extraction and sequencing

Genomic DNA was extracted in one of three ways. 1) 10 mg of hymenophore tissue from each specimen was homogenized in 2.0 ml screw-cap tubes containing a single 3.0 mm and 8 x 1.5 mm stainless steel beads using a BeadBug™ microtube homogenizer (Sigma-Aldrich, #Z763713) for 120 seconds at a speed setting of 3500 rpm. After physical disruption, DNA was extracted using the Monarch® Genomic DNA Purification Kit (New England Biolabs, Massachusetts; #T3010) with the Monarch® gDNA Tissue Lysis Buffer (#T3011) using double the volume of lysis buffer, one hour lysis incubation at 56 C, and 550 µl of wash buffer during each of the wash steps. 2) an in-house 96-well plate protocol where tissue is physically homogenized, as above, after which 1000 µL of lysis buffer (1% sodium dodecyl sulfate, 10 mM Tris, 10 mM EDTA, 5mM NaCl, 50mM dithiothreitol, pH 8.0) is added. To this solution is added 4 µL of RNAse A (20 mg/mL), the solution is vortexed, and then incubated at 37 C for 10 min. Next, 10 µL of proteinase K (20 mg/mL) is added, the solution vortexed, and then incubated at 56 C overnight on an Eppendorf ThermoMixer® with agitation at 400 rpm. After lysis, the tubes are centrifuged at max speed (17,000 x g) to pellet the cellular debris. 700 µL of supernatant is removed to a new 1.7 mL microcentrifuge tube with hinged cap to which 162.5 µL 3.0 M potassium acetate (pH 5.5) is added. The solution is mixed briefly and then put on ice for five min, followed by a second centrifugation, as above. Avoiding the pellet, the supernatant is removed to a well of a 96-well 10 µM filter plate (Enzymax, #EZ96FTP) set in a 2 mL MASTERBLOCK® collection plate (Grainer, #780271). Filtration is achieved through centrifugation at 1500 x g for 2 min. The flow-through is transferred to a new 1.7 ml microcentrifuge tube with hinged cap and centrifuged, as above. Without disturbing the pellet, the supernatant is removed to a new 2.0 mL microcentrifuge tupe with hinged cap and 1000 µL of binding buffer (5M guanidium hydrochloride, 40% isopropanol) is added and the solution homogenized by pipetting. The binding solution is then transferred to a well of a 96-well long-tip AcroPrep™ Plate (PALL, #8133) that was pre-conditioned by pulling 400 µl Tris-HCl buffer (pH 8.0) through using a vacuum manifold. DNA is bound to the filter by centrifugation at 1500 x g for 2 min or using a vacuum manifold. The filter is washed twice with 700 µl of wash buffer (20% solution of 80mM NaCl, 8mM Tris-HCl, pH 7.5 and 80% ethanol) using centrifugation or vacuum, and then the filter is dried with centrifugation at 1500 x g for 15 min. Residual ethanol is removed by incubating the filter plate at room temperature for 30 min. To elute the DNA from the filter, 50 µl of elution buffer (0.1x Tris-EDTA buffer, pH 8-9) prewarmed to 60 C is added directly to the filter, incubated for 2 min at room temperature, and eluted into a new 2 ml MASTERBLOCK® collection plate with centrifugation at 1500 rpm for 2 min. The elution step is repeated once. 3) a phenol-chloroform DNA extraction protocol where tissue is physically homogenized, as above, and lysed using the Tissue Lysis buffer from the Monarch® Genomic DNA Purification Kit (NEB, #T3010S) with double the volume of lysis buffer and a 1 h incubation at 56 C. Then, total lysate was placed in Phase Lock Gel™ Light tubes (QuantaBio, #2302820) along with an equal volume of OmniPur® Phenol:Chloroform:Isoamyl Alcohol (25:24:1, TE-saturated, pH 8.0) solution (MilliporeSigma, Calibiochem #D05686) and then mixed by gentle inversion for 15 minutes using a fixed speed tube rotator. After mixing, tubes were centrifuged at maximum speed (14,000 x g) for 10 minutes, then the aqueous (top) layer was transferred to a new phase-lock gel tube and the process repeated. DNA precipitation of the aqueous phase was performed by adding 5M NaCl to a final concentration of 0.3M and two volumes of room temperature absolute ethanol, inverting the tubes 20x for thorough mixing followed by an overnight incubation at −20C. DNA was pelleted by centrifugation at 14,000 x g for 5 min, washed twice with freshly prepared, ice cold 70% ethanol, air-dried for 15 min at room temperature, and then resuspended in 150 µl of Elution Buffer from the Monarch® Genomic DNA kit.

DNA extract quality was assessed for quality using a NanoDrop 1000 (Thermo Scientific) and fragment integrity using agarose gel electrophoresis. Genomic DNAs were sequenced using a combination of paired-end sequencing on the Illumina MiSeq, HiSeq, and Novaseq sequencing platforms (SUPPLEMENTARY TABLE 2). All raw reads and whole genome assemblies are deposited in the Short Read Archive (Bioproject#PRJNA1022813).

### Genome assembly, ortholog extraction and phylogenetic analysis

Raw sequencing reads were quality-filtered and adapter-trimmed using fastP v0.20.1 (Chen et al. 2018) with default settings. Genome assemblies were produced from quality-filtered reads using SPAdes v3.15.0 (Bankevich et al. 2012) with five k-mer values (k=77,85,99,111,127). From each genome, we identified 1764 highly conserved single copy orthologs using BUSCO with the “basidiomycota odb 10” dataset. Orthogroups that were present in less than 75% of taxa and taxa with less than 20% ortholog recovery were removed. Retained orthologs were aligned using MAFFT v7.397 (Katoh, Rozewicki, and Yamada 2017) with the “L-INS-I” algorithm, and maximum-likelihood gene-trees were inferred using IQ-TREE v2.0.3 (Minh et al. 2020) with automatic model selection in ModelFinder (Kalyaanamoorthy et al. 2017) and ultrafast bootstrapping (BS, (Hoang et al. 2018)) with 1000 replicates. A summary coalescent species tree was constructed from the resulting gene trees using hybrid-ASTRAL implemented in ASTER (v1.15) (Zhang and Mirarab 2022). Branch lengths in substitutions/site were estimated under maximum likelihood on the species tree using the “-te” option in IQ-TREE, with a partitioned concatenated alignment of all BUSCO genes used in species tree construction.

### Gene tree comparison

To evaluate discordance, individual gene trees were compared using six metrics calculated in SortaDate (average bootstrap support, clocklike branch lengths, tree length; Smith et al. 2018) and the R package ‘TreeDist’ (generalized Robinson-Foulds metrics; Smith 2020, 2022). In addition to data matrix summaries (number of taxa, alignment length), Pearson’s correlations were calculated to determine relationships between metrics.

### Divergence dating

A timetree was inferred by applying the RelTime method (Tamura et al. 2012, 2018) conducted in MEGA11 (Stecher et al. 2020, Tamura et al. 2021) to the species tree with ML-estimated branch lengths. To reduce computational burden, time-calibrated branch lengths were calculated using the Maximum Likelihood (ML) method and the General Time Reversible substitution model (Nei and Kumar 2000) from two sets of 100 genes: 1) the top 100 genes with well-supported clock-like trees determined using SortaDate (Smith et al. 2018) and 2) the top 100 genes with the smallest generalized Robinson-Foulds (‘gRF’) distances to the species tree calculated using the R package ‘TreeDist.’ The timetree was computed using two sets of calibration constraints. The first included two calibrations: 1) a secondary calibration for the stem age of the Boletaceae from 50-150 mya (Varga et al. 2019, Wu et al. 2022) and 2) a secondary calibration for the stem age of *Boletus edulis* from 5-13 mya (Tremble et al. 2022). The second set included the former two calibrations plus four of the five internal calibrations using the highly supported core shifts from Varga et al. (2019). Because many of the clades in Varga et al. were incongruent with our topology, calibrations were selected using the most inclusive node, except for *Aureoboletus* which could not be reconciled with our results. The Tao et al. (2020) method was used to set minimum and maximum time boundaries on nodes for which calibration densities were provided, and to compute confidence intervals. Outgroup node ages were not estimated because the RelTime method uses evolutionary rates from the ingroup to calculate divergence times and does not assume that evolutionary rates in the ingroup clade apply to the outgroup.

### Ancestral state reconstruction

Morphological/macrochemical traits were coded according to original descriptions and verified with microscopic analysis when traits were ambiguous. Four traits were scored as binary or multistate characters: 1) hymenophore anatomy (straight tubes = 0, tubular with cross walls = 1, lamellate = 2, not applicable = 3), 2) color changes from damage (none = 0, blue = 1, black/brown = 2, not applicable = 3), and sporocarp morphology (pileate-stipitate with exposed hymenophore = 0, secotioid = 1, gasteroid = 2), 4) spore ornamentation (smooth = 0, not smooth = 1) (SUPPLEMENTARY TABLE 4). The state of hymenophore anatomy defined as “tubular with cross walls” refers to hymenophores that have tubes at two or more lengths, giving the appearance of a primary long tube with shorter internal cross-walls. A “not applicable” category was scored for hymenophore anatomy and color changes from damage to accommodate secotioid/gasteroid taxa and taxa with unknown changes, respectively. An alternative coding scheme for sporocarp morphology was also used in an attempt to disentangle the transition between pileate-stipitate and gasteroid morphologies from the loss of ballistospory. In this alternative coding scheme, two binary traits were scored: stipe (present = 0, absent = 1) and ballistospory (no = 0, yes =1). Ancestral state reconstruction of the root node of the Boletaceae was implemented with BayesTraits V4.0.0, using the MCMC approach over 1,100,000 iterations, with a “burn-in” of 100,000 iterations (Mead and Pagel 2022). Model convergence was assessed with the program Tracer (v1.7.1, Rambaut et al. 2018), and determined as an effective sample size (ESS) of >300 for all variables.

### Ancestral Range Reconstruction

Numerous analytical methods for reconstructing historical biogeography exist, accounting for processes such as vicariance, dispersal, and cladogenesis (Ronquist 1994; Ree et al. 2005; Landis et al. 2013). To account for these macroevolutionary processes in our ancestral state reconstruction in the Boletaceae, we utilized BioGeography with Bayesian (and likelihood) Evolutionary Analysis with R Scripts (“BioGeoBEARS”; Matzke 2018). Samples were coded in two ways: 1) Paleotropical (consisting of Africa and tropical Asia), Neotropical (South and Central America), South Temperate (temperate Australia and New Zealand), or North Temperate (North America, Europe, northern temperate Asia), and 2) by these floristic regions: Holarctic including Central America, Neotropical, Chilean-Patagonian, African, Indo-Malesian, Australian, and Novo-Zealandic (Liu et al 2023) (SUPPLEMENTARY TABLE 3). Central America was combined with the Holarctic region as Central American ECM fungi are mostly derived from North American ancestors (Halling 1996). The most likely model was chosen according to AIC and weighted AIC score calculated in BioGeoBEARS.

## RESULTS

### DNA sequencing, genome assembly, and ortholog extraction

Whole genome sequencing of 418 specimens resulted in 13,794,532 paired-end reads per specimen on average (SUPPLEMENTAL TABLE 2). On average, genome assemblies possessed an assembly N50 of 12.9 Kbp (thousand base-pairs), total assembly length of 61.6 Mbp (million base-pairs), 53,972 scaffolds, and a BUSCO score of 74.7%. 34 of 418 specimens possessed BUSCO scores less than 20%, and 175 specimens possessed BUSCO scores greater than 90% (SUPPLEMENTARY TABLE 2). After removing specimens with poor BUSCO recovery, our final dataset included 383 Boletaceae specimens, three outgroup taxa, and 1461 single-copy orthologs.

### Phylogenetic analysis

The summary coalescent tree resolved most nodes with strong support (FIG. 2). Many of the groups recovered are consistent with previous studies but now with statistical support (Dentinger et al. 2010, Nuhn et al. 2013, Wu et al. 2014). We recognized the six subfamilies following previous authors (Wu et al. 2014), which led us to formally recognize two new subfamilies (Tremble et al. 2023). Many of the currently accepted genera that are not mono- or oligo-typic are polyphyletic. One notable pattern is the phylogenetic placements of the endemic Chilean taxa, all of which were recovered as ancient lineages of similar age within four of the subfamilies: *Gastroboletus valdivianus* in Xerocomoidae, *Boletus loyita* in Austroboletoidae, *Boletus loyo* in Suillelloidae, and *Boletus putidus* in Boletoidae.

**Figure 2.**
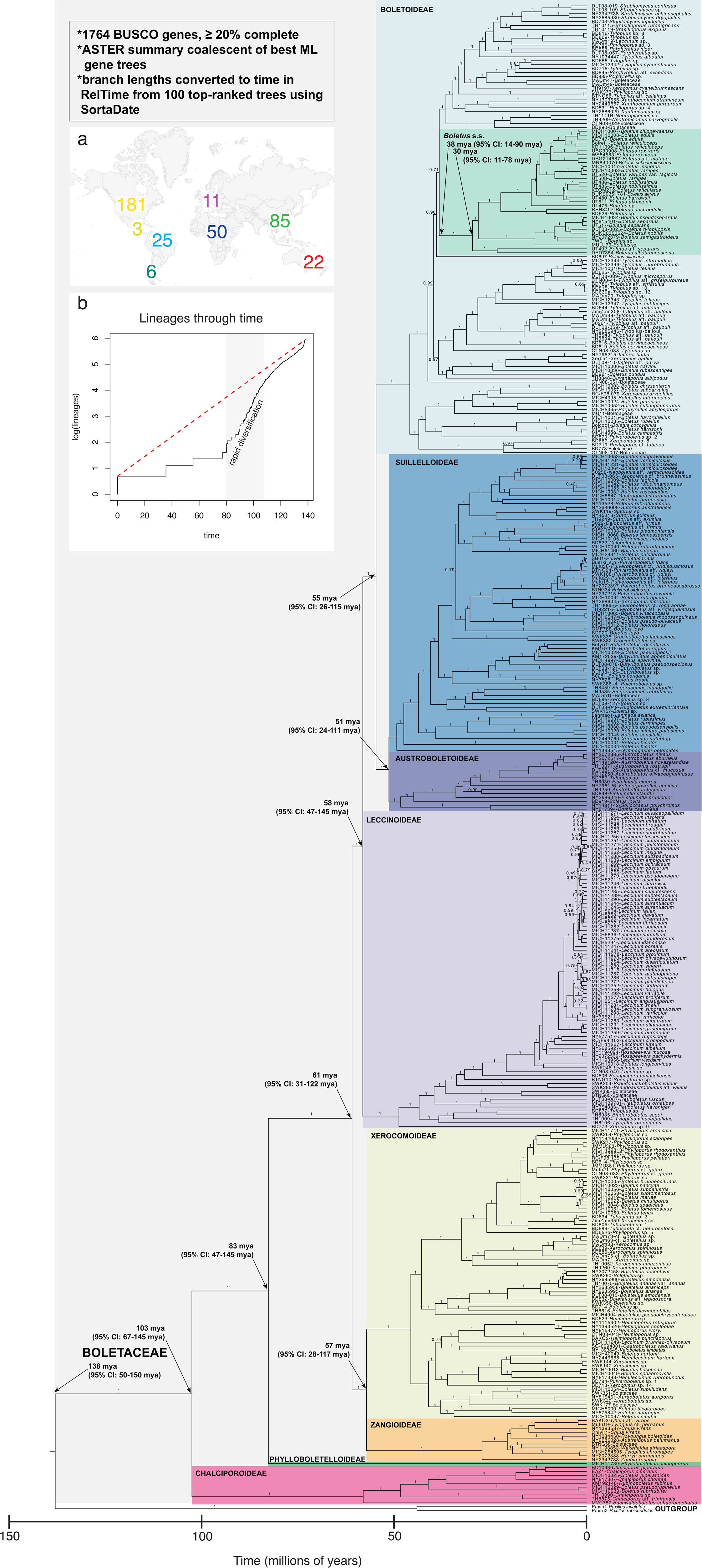
Time-calibrated phylogeny of Boletaceae using 1764 BUSCO genes. Topology is a summary coalescent of individual best ML gene trees using astral-hybrid. Numbers on branches are quartet probabilities. Branch lengths were converted to time using the top 100 best gene trees estimated using SortaDate in RelTime. Inset a) map of specimen origins with numbers of specimens from each geographic area. Inset b) lineages-through-time plot calculated with the “ltt.plot” function in the R package “ape.” The dashed line represents a constant birth-death rate. The shaded box indicates a period of significant divergent increase from a constant birth-death rate indicative of a rapid radiation.

### Gene tree comparison

Average bootstrap support had the highest positive correlation with the number of taxa present (Pearson’s coefficient = 0.76), and weak to moderate negative correlations with alignment length (Pearson’s coefficient = −0.17), clocklike branch lengths (Pearson’s coefficient = −0.13), and total tree length (Pearson’s coefficient = −0.20). Generalized Robinson-Foulds distances were weakly to moderately negatively correlated with number of taxa (Pearson’s coefficient = −0.33), total length (Pearson’s coefficient = −0.30), and clocklike branch lengths (Pearson’s coefficient = - 0.13), and weakly to moderately positively correlated with alignment length (Pearson’s coefficient= 0.29) and average bootstrap support (Pearson’s coefficient = 0.17). Clocklike branch lengths and total tree length were weakly positively correlated (Pearson’s coefficient= 0.15).

### Divergence dating

Using the two- and six-calibration sets the following ages were estimated (FIGS. 2, 3). Stem ages for the Boletaceae were estimated at 138-139 mya and 77 mya. The crown age of the Boletaceae and origin of Chalciporoideae was estimated at 103-105 mya and 63 mya (49-77 mya). The stem age of the Phylloboletelloideae was estimated at 83-87 mya and 58 mya (49-77 mya)., The radiation of the remaining subfamilies was estimated to have occurred between 61 and 51 mya. The origin of *Boletus* sensu stricto (i.e. “true porcini”) was estimated at 38 mya and 35 mya (29-42 mya), and its diversification was estimated at 29-30 mya and 26 mya (20-34 mya).

**Figure 3.**
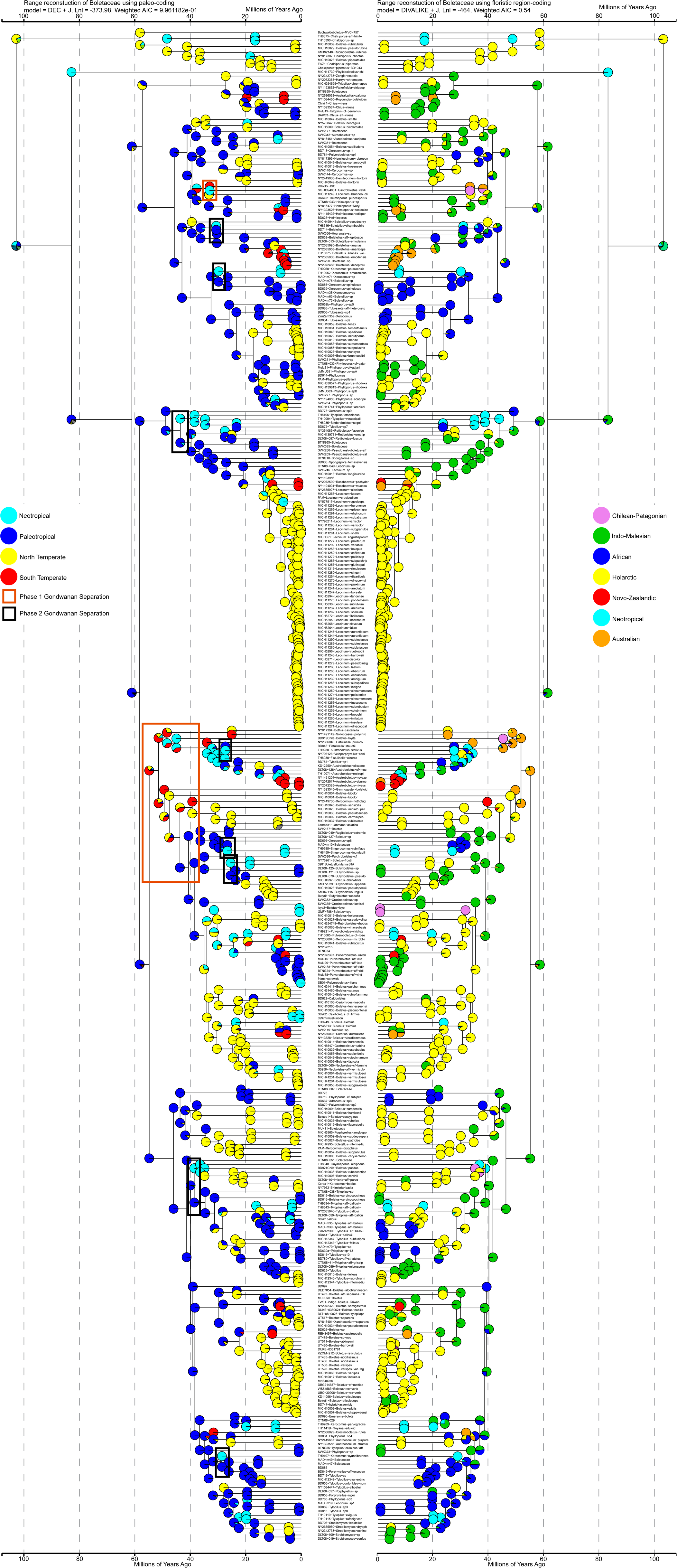
Biogeographic reconstruction using BioGeoBEARS. Left-hand tree depicts 4-state coding scheme (light blue=Neotropical, blue=Paleotropical, yellow=North Temperate, red=South Temperate) and right-hand tree depicts floristic region coding scheme (pink=Chilean-Patagonian, green=Indo-Malesian, blue=African, yellow=Holarctic, red=Novo-Zealandic, light blue=Neotropical, orange=Australian). Pie charts indicate the proportional likelihood of each state at a node. Red and black boxes indicate Phase I and Phase II Gondwanan diversification events, respectively.

### Ancestral state reconstruction

Ancestral state reconstruction with Bayestraits achieved strong convergence (all variables with ESS > 300) and recovered the most likely ancestor of the Boletaceae to have a pileate-stipitate, lamellate sporocarp with ornamented basidiospores. The highest transition rates were observed in changes from secotioid to gasteroid and from secotioid to pileate-stipitate morphologies. Secotioid morphology appeared to be a relatively transient evolutionary state (TABLE 1). High transition rates were also found to occur from gasteroid to pileate-stipitate morphology, supporting evidence for the reversibility of gasteromycetization.

**Table 1.** Morphological transition rates estimated with Bayestraits.

| **Trait** | **Transition** | **Mean rate** | **Median rate** |
| --- | --- | --- | --- |
| **Ballistospory** |  |  |  |
|  | Yes to No | 1.2303 | 1.0565 |
|  | No to Yes | 39.8088 | 35.7478 |
| **Spore Ornamentation** |  |  |  |
|  | Smooth to Not Smooth | 0.7747 | 0.7498 |
|  | Not Smooth to Smooth | 1.741 | 1.464 |
| **Stipe** |  |  |  |
|  | Absent to Present | 33.685 | 27.67 |
|  | Present to Absent | 0.8415 | 0.6788 |
| **Sporocarp Habit** |  |  |  |
|  | Secoitiod to Pileate-stipitate | 51.64 | 51.47 |
|  | Pileate-stipitate to Secoitiod | 0.8178 | 0.7172 |
|  | Secoitiod to Gasteroid | 53.44 | 53.90 |
|  | Gasteroid to Secoitiod | 34.63 | 31.05 |
|  | Pileate-stipitate to Gasteroid | 0.8393 | 0.6749 |
|  | Gasteroid to Pileate-stipitate | 40.78 | 36.85 |
| **Hymenophore Bruising** |  |  |  |
|  | None to Blue | 1.3515 | 1.2632 |
|  | None to Black | 0.6921 | 0.6921 |
|  | Blue to None | 5.3730 | 5.3540 |
|  | Blue to Black | 0.1732 | 0.1205 |
|  | Black to None | 3.2329 | 2.7385 |
|  | Black to Blue | 1.0294 | 0.7281 |
| **Hymenophore Anatomy** |  |  |  |
|  | Straight to Cross | 0.6398 | 0.6092 |
|  | Straight to Lamellete | 0.2054 | 0.1715 |
|  | Cross to Straight | 3.694 | 2.960 |
|  | Cross to Lamellate | 2.4131 | 1.6755 |
|  | Lamellate to Straight | 5.483 | 5.004 |
|  | Lamellate to Cross | 1.858 | 1.275 |

### Ancestral range reconstruction

Ancestral distribution reconstruction recovered a likely Paleotropical origin of the Boletaceae (DEC+J model chosen with lowest AIC and AICc for both coding sets), with two major descendant radiations originating in the Paleotropics (Africa and Asia) and Neotropics (FIG. 3). In addition, we found evidence for multiple diversification events spurred by the separation of Gondawana (FIG. 3). Gondwanan separation occurred in two predominant phases: Phase 1, which involved the separation of Southern South America, Southern Africa, Australia-Antarctica, and Madagascar-India, beginning approximately 180 mya and largely completed by 120 mya (Jokat et al. 2003) and Phase 2, involving the separation of South America and Africa, which was completed 80 mya (Reguero and Goin 2021). At the split between the Austroboletoidae and Suillelloidae (FIG. 3), we estimated a putative Phase 1 Gondwanan separation to have occurred 150-120 mya, which led to rapid formation of South Temperate, Neotropical, and Paleotropical lineages. Later, around 90-70 mya, at least five putative Phase 2 separation events occurred, splitting the Paleotropical and Neotropical lineages.

In our four category paleo-region coding set, the Boletaceae ancestor was equally likely to be Neotropical or Paleotropical. However, the subsequent node that leads to the rest of the Boletaceae (excluding Phyllobolletoidae and Chalciporoideae) was well-supported as Paleotropical, as were all immediate descendent nodes. Our coding of the Chalciporoideae and the single *Phylloboletellus* specimen likely had a strong influence on deep-node ancestral range reconstructions. The backbone nodes of the Boletaceae excluding Phylloboletoidae and Chalciporoideae were estimated as Asian in origin, corroborating the four-category analysis, though with less confidence. Migrations between phytogeographic regions were dominated by dispersals between the Indo-Malesian and Holarctic regions (FIGS. 3,4).

**Figure 4.**
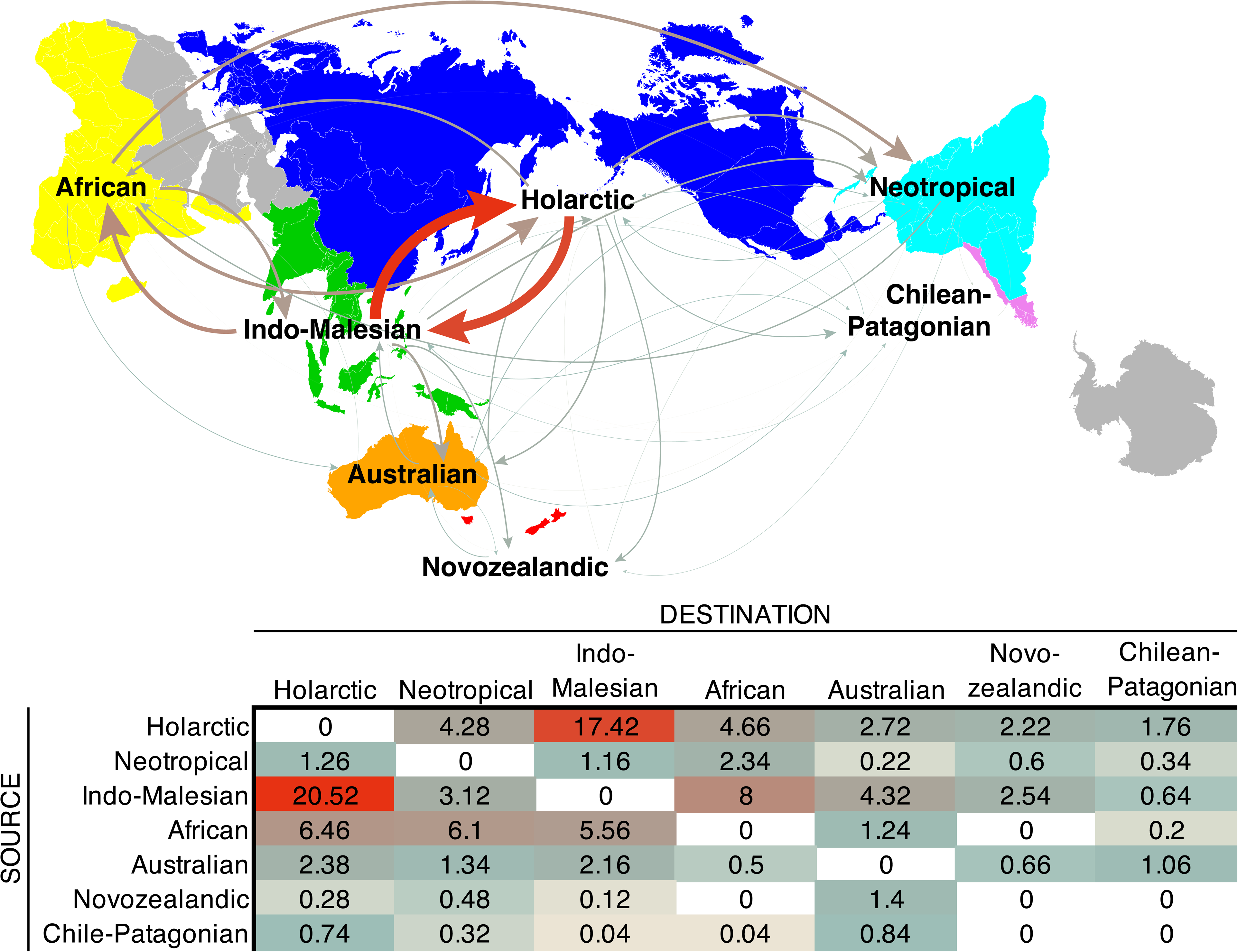
Dispersal events and rates inferred from BioGeoBEARS. Top: Map depicting phytogeographic zones that are colored and labeled on the map. Map was ren-dered using the ‘imago’ R code (https://github.com/hrbrmstr/imago) to reproduce the AuthaGraph world map projection (http://www.authagraph.com/top/?lang=en). Curved arrows indicate inferred directional dispersal events and are colored by rate values following the table (Bottom). The stroke weight of the arrows has been scaled to percent of the maximum rate value following the values in the table. Bottom: Table of dispersal rates inferred under a DIVAlike+j model in BioGeoBEARS. Source regions are at left and destination regions are along the top. Values are colored along a scale from cool to warm (red being maximum).

**Figure 5.**
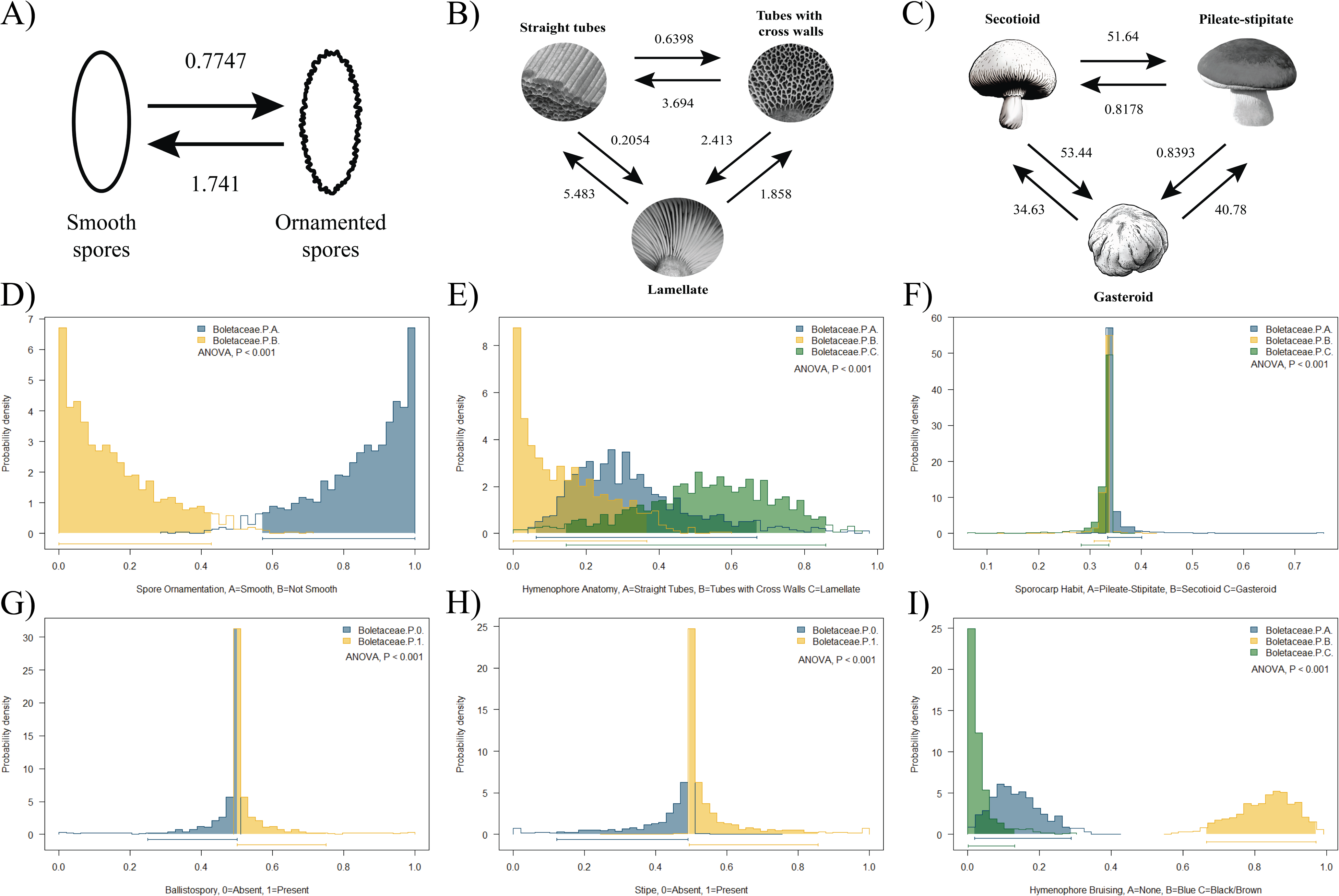
Distribution of likelihood frequencies of morphological traits. A-C) Illustrations of coded traits and inferred transition rates. Arrows indicate direction of transition with stroke weights scaled to maximum rate value. D-I) Frequency distributions of state probabilities for six trait codings. Traits and codes are labeled on the x-axis.

## DISCUSSION

The fully resolved phylogeny supports the recognition of eight subfamilies, including the newly defined Phylloboletelloideae and Suillelloideae (Tremble et al. 2023). The subfamilial relationships were resolved for the first time. The Chalciporoideae was recovered as the earliest diverging group in the Boletaceae, a relationship previously noted by Wu et al. (2014). The enigmatic *P. chloephorus* (Singer and Diglio 1952) was the next lineage to branch off before the radiation that gave rise to the six additional subfamilies. Previous studies have placed *Pseudoboletus parasiticus* in a position similar to that of *P. chloephorus* in our study (Nuhn et al. 2013, Wu et al. 2014, Sato and Toju 2019, Caiafa and Smith 2022). We were, however, unable to include a representative of *P. parasiticus* in our study so cannot corroborate its putative phylogenetic position.

The tree topology has intriguing implications for the role of ecological transitions in Boletaceae diversification. Members of the earliest-diverging Chalciporoideae species can be facultatively ECM, saprotrophic or mycoparasitic (Caiafa and Smith 2022) and *P. chloephorus* may not be ECM given its occurrence in non-ectotrophic forests (Singer and Diglio 1952). Moreover, *P. parasiticus* and other *Pseudoboletus* spp. produce sporocarps directly attached to gasteroid *Scleroderma* and *Astraeus* and are assumed to be mycoparasites (Raidl 1997; Binder and Hibbet, 2006; Nuhn et al. 2013). Altogether, the basal position of these early diverging groups suggests that the ancestor of the Boletaceae was likely saprotrophic and not ECM. This possibility is in line with the results of Sato and Toju (2019) which indicated that the ECM habit emerged with the origin of the six derived Boletaceae subfamilies. Genomic changes coinciding with the emergence of an obligate ECM habit further support the view that this nutritional shift has profoundly impacted Boletaceae (Wu et al. 2022).

Many taxonomic changes in the Boletaceae have been proposed in recent years. In particular, new genera have been erected for phylogenetically unresolved lineages. Many of these new genera are mono- or oligo-typic (composed of one or few species) (e.g. Castellano et al. 2016; Chakraborty et al. 2015; Henkel et al. 2016; Halling et al. 2023). This proliferation of new genera has made it difficult or impossible to recognize inclusive groupings without requiring knowledge of the species. On the bright side, the strong nodal support throughout our phylogeny sets the stage for a new, comprehensive and stable generic-level taxonomy. We will address this in subsequent works when all recently described genera are represented.

Our biogeographic and divergence dating analyses support a Gondwanan origin of the Boletaceae. Subsequent divergence was likely facilitated by continental drift-based vicariance events and possible long-distance dispersals. Other recent studies have shown that lineages of ECM fungi originated in Gondwana or more recently in paleotropical regions (Hosaka et al. 2008, Matheny et al. 2009, Dentinger et al. 2010, Ryberg and Matheny 2011, Kennedy et al. 2012, Sanchez-Ramirez et al. 2015, Han et al. 2018; Hackel et al. 2022; Codjia et al. 2023). Phytogeographic endemism was implied with our biogeographic reconstruction. The strongest migrations occurred recently over the past 50 my between the Indo-Malesian and Holarctic regions. We acknowledge the difficulty to determining origins and dispersal events in the absence of fossils or other corroborating evidence. Nonetheless our study and others suggest that vicariance may have played a strong role in the distribution of ECM fungal taxa, despite the long-distance dispersal capacity of airborne spores (Matheny et al. 2009, Peay et al. 2010, Peay and Matheny 2016). Conversely, pure vicariance cannot explain the close phylogenetic relationships seen between distantly disjunct taxa. Long-distance dispersals may have occurred, albeit rarely. While long-distance dispersal is demonstrably possible in the Boletaceae and other ECM lineages the likelihood of its frequent occurrence is low (Geml et al. 2012; Hackel et al. 2022; Tremble et al. 2022). Most basidiospores do not travel far from the parental sporocarp (Galante et al. 2011), and the probability of two airborne basidiospores landing in close-enough proximity to mate is negatively correlated with increasing distance from sporocarps (Peay et al. 2012; Golan and Pringle 2017). Such improbabilities notwithstanding, our biogeographic patterns are consistent with episodic long-distance dispersal, possibly by aerial dispersal of basidiospores, spores vectored by migrating animals (e.g., Elliot et al. 2019) or somatic mycelia on rafting vegetation (Thiel et al. 2005).

Our biogeographic reconstructions are consistent with the “Southern Route to Asia” hypothesis (Wilf et al. 2019). This idea proposes that ECM Fagaceae and their symbiotic fungi originated in Gondwana and were carried on Australia north to Asia. In this scenario the Gondwanan ECM habitat tracked climatic niches on the desertifying continent northward as the Australian plate collided with the Pacific plate. A relictual ECM community remained in a newly isolated New Guinea and subsequently spread northwest along the montane Australasian archipelago, followed by dispersal into continental Asia. Many of the dispersal events we found between Indo-Malesia and other regions, especially the Holarctic, are inferred within the last 20 my, coincident with the late-Oligocene collision of Australia with the Pacific plate (Hall 2011). As suggested by Halling et al. (2011) recent Boletaceae migrations likely occurred across the Australasian archipelago and are corroborated by our inferred recent regional dispersal events.

Biogeographic reconstructions are highly sensitive to taxon sampling and our dataset is not immune to equivocal reconstructions. For example, in both coding schemes the ancestral node of the Chalciporoideae had the highest probabilities of a North Temperate and North American origin, respectively. However, with no Chalciporoideae samples from Asia, Africa or Australia/New Zealand in our study, their potential impacts on the reconstruction are unknown. Such sampling gaps notwithstanding, we have the most geographically comprehensive sampling for Boletaceae ever compiled and provide the first opportunity to examine global-scale biogeographic patterns. Insights into the evolution of the Boletaceae are revealed for the first time, despite mild uncertainty at a minority of nodes.

The evolutionary origins of distinctive regional Boletaceae assemblages have long been a mystery (Horak 1977). For example, the endemic Boletaceae of Chile and Argentina have not been included in previous phylogenetic studies, and their morphology-based affinities have been inconclusive (Horak 1977). The recovery of several Chilean species as ancient lineages in four of the subfamilies implies that they have survived in isolation without speciating for millions of years. The closest relatives of these Chilean boletes occurred in geographic regions as disjunct as North America, lowland tropical northern South America, and Australia. *Boletus loyita* and *G. valdivianus* were most closely related to extant Australian taxa, suggesting an origin prior to final Gondwanan disarticulation (Reguero and Goin 2021). Close relationships between southern Gondwanan Australian and southern South American taxa have been documented elsewhere (Feng et al. 2017). In all likelihood Chilean boletes arose in Gondwana, separated from their sister lineages during Gondwanan disarticulation, and underwent no subsequent speciation for tens of millions of years.

Ancestral range reconstruction recovered an Asian origin of the core, “true porcini” genus *Boletus* s. str. as previously suggested (Feng et al. 2012). However, we cannot entirely rule out an African origin. The Central African endemic *Boletus alliaceus* was recovered here as a sister taxon to *Boletus* s. str., and a similar relationship was found for the recently described *Paxilloboletus africanus* (Badou et al. 2022). Furthermore, we estimated the origin of *Boletus* s. str to be 40 mya, which may indicate why the sister lineages to *Boletus* s. str. are endemic to Africa. India had separated from Africa and Madagascar ∼120 mya (Reguero and Goin 2021), and at 40 mya was already colliding with Asia (Aitchison et al. 2007; Hu et al. 2016). If *B. alliaceus* and *P. africanus* are indeed sister lineages of *Boletus* s. str., then the arrival and subsequent diversification of true porcini in Asia must have been a dispersal event, because the separation of India from mainland Africa ∼180-170 mya (Hankel 1994) occurred long before our estimated age of the *Boletus* s.s. ancestor (∼40 mya). Even if a more recent ancestor existed in Madagascar or the Seychelles, the separation of India from these landmasses at ∼90 mya (Storey et al. 1995) and ∼64 mya (Norton and Sclater 1979), respectively, is still much older than our current age estimates for true porcini. Furthermore, most or all ECM fungi in Madagascar appear to have arrived on the island through dispersal its separation from Africa (Rivas-Ferreiro et al. 2023), so dispersal is the most plausible mechanism unless ancient Malagasy relict taxa are discovered. In the current study currently undescribed species of *Boletus* s. str. were recovered from Taiwan, Malaysian Borneo, and the Gulf Coast of the US, indicating that much more diversity exists in the genus. To sort out the origins and full diversity of *Boletus* s. str. more mycological exploration and whole genome sequencing are needed. In particular, discovery and analysis of true porcini basal lineages from India and Africa could shed further light on the origin of this charismatic group.

Divergence dating estimated the origin of the Boletaceae at 138-139 mya, and ancestral range reconstructions suggested it may be even older. Ranking genes with different metrics had little impact on divergence date estimates. However, the two calibrations sets gave very different estimates for most nodes. The estimated origin dates of the Boletaceae using the Varga et al. (2019) calibrations were almost half those of the two-calibration set. It is difficult to interpret these wildly different divergence dates given the lack of fossil evidence. However, the dates estimated using the Varga et al. (2019) calibrations are suspect due to extensive topological incongruence of their phylogenetic trees with ours. Our older divergence estimate corroborates the results of He et al. (2020) and our internal dates correspond with other results, such as the ∼48 mya origin of the *Strobilomyces* group (Han et al. 2018). Our older divergence estimate is also in line with the origin of ECM Pinaceae in the early Cretaceous (Brundrett and Tedersoo 2018). Therefore, we consider the older estimate to be more accurate.

In our ancestral range reconstruction analysis, we found evidence of multiple diversification events that may have been initiated by Gondwanan breakup. The first phase of the Gondwanan separation postulated by Jokat et al. (2003) correlates well with our estimated origin of the Boletaceae and indicates that the family was diverse and widely distributed by 120 mya, substantially older than the estimated age from our divergence dating analysis but within the 95% confidence interval. Our dates are at best coarse estimates based on fossil-free secondary calibrations. However, the phylogenetic pattern of vicariance that parallels the breakup of Gondwana is compelling and offers corroborating evidence that our estimated ages may in fact be too young. In any case, our divergence estimates suggest that the Boletaceae originated and diversified within the early to late Cretaceous period. During this time global climate was warm and wet (Hay and Floegal 2012), gymnosperms and subsequently angiosperms diversified (Crisp and Cook 2011), and the supercontinental land masses broke apart (Jokat et al. 2003).

The Boletaceae mostly consists of species that form ECM assocations, but emerging evidence suggests the ancestor may have had mycoparasitic capacity. The gain of obligate ECM ecology in the six most-derived subfamilies likely occurred after their divergence from the Phylloboletelloidae. *Phylloboletellus chloephorus* is from non-ECM dominated habitats. The Chalciporoideae have not been definitively shown to form ECM associations but do have saprotrophic or mycoparasitic capacities (Caiafa et al. 2022). While mycoparasitic *Pseudoboletus* species have also been recovered as early-diverging Boletaceae lineages (Nuhn et al. 2013, Sato and Toju 2019, Cortes-Perez et al. 2023) and may have close affinities with lamellate *Phylloboletellus*, we were not able to evaluate *Pseudoboletus* in this study. A more thorough investigation of the ecology of early-diverging Boletaceae is needed to test this “mycoparasitic origin” hypothesis.

A longest standing debate among Boletaceae systematists has centered on the utility of morphological traits for defining natural genera. Basidiospore color, ornamentation, and hymenophore arrangement were long emphasized in this respect. These features have been used as evidence for dividing the Boletaceae into multiple genera (e.g., Singer 1945a,b; 1947; Pegler and Young 1981) or treating nearly all Boletaceae as a single genus (Corner 1972). More recently, hyphal anatomies of sporocarp structures and pigment chemistry have been emphasized (e.g. Binder et al. 2002; Šutara 2005). Yet, despite the morphological and chemical variability in the Boletaceae, the family is typified by the ‘bolete’ macromorphology of fleshy, pileate-stipitate sporocarps with tubular hymenophores. In addition, type of basidiospore ornamentation has been long considered to be a genus-unifying trait in genera such as *Strobilomyces*, *Boletellus*, and *Austroboletus* (Berkeley 1851, Murrill 1909, Corner 1972, Pegler and Young 1981, Wolfe 1980). Despite such traditional views, our ancestral state reconstructions suggest that the ancestor of the Boletaceae had a pileate-stipitate sporocarp with a lamellate hymenophore and ornamented basidiospores. This likely resulted from the basal phylogenetic position of the lamellate *P. chloephorus* (Singer and Diglio 1952).

The evolution of sequestrate morphologies has long been thought to be irreversible (Thiers 1984). The transition from pileate-stipitate to secotioid to gasteroid morphology involves enclosure of the hymenophore and loss of ballistospory, which are unlikely to be regained once lost (Hibbett et al. 1997, Hibbett 2004, Sanchez-Garcia et al. 2020). However, prior studies have not fully rejected the hypothesis that gasteromycetization is irreversible (Hibbett 2004; Wilson et al. 2011; Sanchez-Garcia et al. 2020). Our ancestral state reconstructions under two coding schemes strongly supported transition to and from gasteroid forms. This suggests that gains or losses of ballistospory may also be reversible conditions. In our analyses, transitions between pileate-stipitate with exposed hymenophore, secotioid, and gasteroid forms suggest that the secotioid condition is intermediate, as indicated by the equal rates of transition to gasteroid and pileate-stipitate forms. Moreover, inferred transition rates from pileate-stipitate to both secotioid and gasteroid forms were almost zero, suggesting that the secotioid condition is evolutionarily unstable. This result contrasts with the stability of *Cortinarius* secotioid taxa suggested by Peinter et al. (2001). A plausible hypothesis is that a membranous partial veil that covers the hymenophore at early stages of sporocarp development may predispose it to gasteromycetization. However, although we did not test this explicitly, based on the phylogenetic distribution of taxa with membranous veils (e.g., *Pulveroboletus*, *Veloporphyrellus*), it is clear that the membranous partial veil is a convergent trait not closely associated with secotioid or gasteroid morphologies. A possible exception to this is *Veloboletus limbatus*, which is most closely related to the gasteroid *Gastroboletus valdivianus*, although their common ancestor was estimated at ∼40 mya.

Transitions between hymenophore organization parallels the sporocarp morphologies. The “tubular with crosswalls” hymenophore condition appears to be intermediate between lamellate and tubulate forms, with transitions to tubes being 1.3-5.8 times greater than the opposite transitions. However, the greatest transition rate was from lamellate directly to tubulate forms, indicating that an intermediate morphology may not be necessary or, like the secotioid condition, may be evolutionarily unstable. Our data provide compelling evidence that lamellate hymenophores, color changes, and gasteromycetization have evolved multiple times in the Boletaceae and are reversible.

## SUMMARY

The Boletaceae underwent a rapid radiation and subsequent long period of phylogenetic instability. These issues had long prevented accurate assessment and analysis of trait evolution and conclusive generic-level taxonomic frameworks in the family. Previous molecular phylogenetic studies were based on a limited taxon sampling and a few loci and generated trees with many short branches and little deep-node support. This study provides the first Boletaceae phylogeny with strong support at deep nodes, based on a massive dataset of 1461 single copy genes from 383 genomes sampled from taxonomically and geographically comprehensive specimens. Our analyses indicated that the Boletaceae likely arose before Gondwanan breakup and that present-day distributions are partly due to vicariance. Long-distance dispersal could not be ruled out for some current distributions. Morphological traits resulted from convergence and frequent reversals, which, along with rapid radiation, have long confounded attempts to achieve a natural intrafamilial classification. This study provides new genomic data and a solid phylogenetic framework that will enable a renewed foundational taxonomy as well as deeper analysis of trait evolution.

## AKNOWLEDGEMENTS

This work was supported by NSF-DEB awards DEB-2114785 to BD and DEB-0918591 to TWH and DEB-1556338 to TWH and BD; National Geographic Society’s Committee for Research and Exploration grants (6679-99, 7435-03, and 8481-08 to TWH). JMM was supported by grants from the ROM Governors and the Natural Sciences and Engineering Research Council of Canada, Discovery Program. We would like to thank Dr. Roy Halling for the numerous specimens provided.

## CONFLICTS OF INTEREST

The authors declare no conflict of interest, financial or otherwise.

## Notes

### Competing Interest Statement

The authors have declared no competing interest.

### Summary of Updates

Supplemental table 1 has been updated to include additional voucher information and the manuscript has been slightly revised.

